# COMICS: A pipeline for the composite identification of selection across multiple genomic scans using Invariant Coordinate Selection in R

**DOI:** 10.1101/587469

**Authors:** Joel T. Nelson, Omar E. Cornejo

## Abstract

Identifying loci that are under selection versus those that are evolving neutrally is a common challenge in evolutionary genetics. Moreover, with the increase in sequence data, genomic studies have begun to incorporate the use of multiple methods to identify candidate loci under selection. Composite methods are usually implemented to transform the data into a multi-dimensional scatter where outliers are identified using a distance metric, the most common being Mahalanobis distance. However, studies have shown that the power of Mahalanobis distance reduces as the number of dimensions increases. Because the number of methods for detecting selection continue to grow, this is an undesirable feature of Mahalanobis distance. Other composite methods such as invariant coordinate selection (ICS) have proven to be a robust method for identifying outliers in multi-dimensional space; though, this method has not been implemented for genomic data. Here we use simulated genomic data to test the performance of ICS in identifying outlier loci from multiple selection scans and compare the results to the performance of Mahalanobis distances. We show that the ICS outperformed Mahalanobis distance in all aspects including false positives, false negatives, and recall. Furthermore, ICS also performed better when identifying loci with weaker selection coefficients. We also introduce a pipeline in a R-Shiny smart wrapper environment that implements the ICS on multiple scans of selection. Importantly, we show that the ICS is a robust method for identifying outliers in multi-dimensional space and recommend its use for studies aimed at identifying loci under selection in the genome.

## Introduction

A main goal of evolutionary genetics and ecological genomics is to identify candidate regions of the genome that show patterns consistent with selection (Beaumont and Balding 2004; Campbell‐Staton et al. 2016; Forester et al. 2016; Martins et al. 2016). Highly differentiated loci are often interpreted as regions of the genome under selection, potentially due to local adaptation of a population to a novel or changing environment (Beaumont and Balding 2004; Campbell‐ Staton et al. 2016; Bekkevold et al. 2016; Rellstab et al. 2016). With the improvement of sequencing methods, the quantity and quality of genetic data continues to increase, allowing a more rigorous analysis of highly differentiated loci and how they are distributed throughout the genome. Despite this increase in genetic data methods to identify putatively selected loci (Gunther and Coop 2013; Lotterhos and Whitlock 2015; Capblancq et al. 2018), identifying candidate loci under selection and differentiating between true outliers and false positives still proves to be a difficult task.

Many selection methods/scans include the analysis of single nucleotide polymorphisms (SNPs), where each SNP is treated as an independent hypothesis when tested against the rest of the genome (Nielsen et al. 2005; Lotterhos and Whitlock 2014; Lotterhos and Whitlock 2015). The increase in SNP data ultimately decreases our ability to differentiate between neutral and selective hypotheses, because of concomitant increase in false positives.. This problem is also present when referring to methods that use discrete windows containing multiple SNPs to detect overall deviations in the pattern of variation in local regions when compared to the rest of the genome (Sabeti et al. 2002; Nielsen et al. 2005; Sabeti et al. 2007; Chen et al. 2010; Zhong et al. 2011; Alachiotis et al. 2012; Pavlidis et al. 2013). To circumvent this issue, some studies have implemented the use of multiple selection methods to identify regions of interest and combine them in a meaningful way (*Anopheles gambiae* 1000 Genomes Consortium 2017; Zueva et al. 2018; Hodel et al. 2018). For example, a common method is to only define genomic regions as under selection if they are identified as outliers by multiple selection methods. Because of the nuances across selection methods and the differences in the assumptions and types of signals, the number of intersections identified may underestimate the true number of regions under selection, increasing the false negative rate and making this approach very conservative. For instance, methods that detect selection using the site frequency spectrum (SFS) identify a selective sweep as an area with higher than expected levels of allele differentiation (Nielsen et al. 2005), while methods using linkage disequilibrium identify selective sweeps as regions with larger than expected associations among SNPs (Alachiotis et al. 2012). When looking at the intersection between these methods, regions under moderate selection near high levels of recombination would most likely be considered neutral.

In lieu of looking at the intersection across multiple selection methods to identify outlier loci, composite methods should be implemented. A composite method includes the use of multiple selection statistics to identify regions of the genome under selection (Verity et al. 2017; Capblancq et al. 2018). In this case, each method/statistic will result in a distribution that is described by its location and dispersion. A multi-dimensional transformation creates a cluster of points, derived from the distributions of the selection/population statistics, where the location is defined as the center of the cluster and the spread of the data are defined as the scatter (Verity et al. 2017; Capblancq et al. 2018; Archimbaud, Nordhausen, Ruiz-Gazen 2018a). Moreover, multidimensional analyses create ample opportunity to take different selection metrics and compare them linearly under one distribution (Verity et al. 2017). In the case of multi-dimensional data, most outliers are identified by a distance metric, where they are expected to have a larger than average distances from the central location of the cluster (Archimbaud, Nordhausen, Ruiz-Gazen 2018a).

A popular composite statistic recently proposed to identify outliers across multiple tests of selection is the Mahalanobis distance, a measure of the number of standard deviations that a given data point is from the multi-dimensional mean (Mahalanobis 1936; Gnanadesikan and Kettenring 1972; De Maesschalck et al. 2000; Verity et al. 2017; Archimbaud, Nordhausen, Ruiz-Gazen 2018a). Though the Mahalanobis distance has been implemented for identifying outlier loci in a multi-dimensional data set, recent studies have shown that an increase in the number of dimensions (the number of metrics estimated within a data set) results in an inflation of false negatives and false positives (Leys et al. 2018; Archimbaud, Nordhausen, Ruiz-Gazen 2018a). Specifically, results from Archimbaud et al. (2018) show that if outliers belong to a reduced dimension space, the probability that the Mahalanobis distance of a neutral locus exceeds the distance of a true outlier is low; however, this is not the case when dimension space is increased. In other words, as the number of selection methods used increases, it becomes more difficult to differentiate between true positives and true-negatives. This phenomenon is known as the “curse of dimensionality,” (Trunk 1979; Bellman, Richard 2013; Bellman, Richard E. 2015).

We propose the use of invariant coordinate selection (ICS) as an alternative method to combine multiple selection scans and identify true outliers in growing multi-dimensional space. Here, we also present a pipeline to easily implement the method. Specifically, ICS is a method for identifying outlier data points in multi-dimensional space with coordinates derived from eigenvalues and eigenvectors (Tyler et al. 2007; Archimbaud, Nordhausen, Ruiz-Gazen 2018a). The ICS is analogous to a principal component analysis (PCA), however, it differs in two ways. First, instead of using one scatter matrix (the covariance matrix commonly used in PCA), ICS adds further constraints by solving inequality conditions on a second matrix with either the third moment (skewness) or the fourth moment (kurtosis) (Tyler et al. 2007; Archimbaud, Aurore, Nordhausen, Ruiz-Gazen 2018a). Second, the invariant components are maintained in parallel while principal components are orthogonal (Tyler et al. 2007; Archimbaud, Aurore, Nordhausen, Ruiz-Gazen 2018a). Once the univariate data are transformed into a multi-dimensional data set, data points with the largest ICS distance (i.e., the Euclidean distance) in the transformed space are deemed as outliers. Importantly, because of the information drawn from the two different scatter matrices, ICS methods are less susceptible to the inflation of distance with an increase in the number of dimensions (Tyler et al. 2007; Archimbaud, Aurore, Nordhausen, Ruiz-Gazen 2018a).

ICS has previously been identified as a robust method for identifying outlier in multidimensional space and has been implemented to detect subgrouping within populations based on genotypic data, however, it has not been implemented for identifying outlier regions based on the input of selection scans. (Archimbaud, A. et al. 2016; Fischer et al. 2017; Archimbaud, Nordhausen, Ruiz-Gazen 2018b). We use simulated genetic data to compare the number of false positives, false negatives, and recall between Mahalanobis and ICS distances. We also introduce an easy to use R-Shiny wrapper specifically designed for performing ICS using genomic data. We call this app COMICS (<u>**C**</u>alling <u>**O**</u>utlier loci from <u>**M**</u>ulti-dimensional data using <u>**I**</u>nvariant <u>**C**</u>oordinate <u>**S**</u>election) and make it available at: https://github.com/JTNelsonWSU/COMICS.

## Results

### The COMICS R-package

COMICS was implemented with the R-Shiny environment, which is an open source R-package that provides a framework for building web applications using R (Chang et al. 2015; Team 2018). Specifically, the Shiny environment allows for the development of a graphical user interface (GUI) that permits the user to operate the ICS proficiently. Using the COMICS package, the user is able to analyze data in several different ways. First, COMICS has the option to observe the data before the multi-dimensional transformation; this feature allows the user to view the results from the different selection scans used to generate the multi-dimensional data in individual plots. COMICS allows for the customization of the end results of the ICS identification of outliers by modifying statistical cutoffs, and specific chromosome intervals. COMICS allows the user to download the multi-dimensional data frame and corresponding figures in csv and pdf formats. The pipeline is user friendly while maintaining flexibility in the implementation of the method. You can find the COMICS app and example files at: https://github.com/JTNelsonWSU/COMICS.

### Data upload and input file format

COMICS requires two input data files. The first file is a multivariate data file containing a list of each genomic position with the corresponding likelihood estimate, or effect size estimate, from each scan of selection used. The first two columns contain the chromosome and physical position along the chromosome (or equivalent). COMICS can handle SNP or window analyses, as long as the windows are consistent across scans of selection; in the latter case, the position will correspond to the midpoint of the window. The physical positions need to be numerically organized relative to only the corresponding chromosome; COMICS is designed to convert all physical positions to the length of the genome.. The second data file is known as the genome configuration file. Specifically, the configuration file is used to define the number of chromosomes (or scaffolds) used, including the total length of each chromosomes in base pairs. The main role of this file is to tell COMICS how large the genome (or assembly) is and the order in which the positions align along the genome; this is taken into account when COMICS generates genome-wide selection scan figures.

### COMICS features and figure generation

In addition to customizing statistical thresholds (e.g., quantiles of interest), COMICS will generate several different figures for both single statistics and ICS distances. For example, two different figures are generated from the multivariate data prior to the analysis. The first is a scatter plot generated by the ggplot and reshape R-packages (Wickham 2007; Wickham 2016), consolidating all genome scans into one panel and color codes them by chromosome (Figure 1). The second figure is a user-defined representation of one particular scan of selection at a time. After analyses with ICS, COMICS will produce two histograms of the log ICS distance for 1) the entire genome and 2) a user defined chromosome. Each of these has a vertical line that defines a cutoff based on the quantile of interest. The main figure that COMICS produces is a genome scan of the logged ICS Euclidean distance. Like the univariate genome scans, the ICS genome scan is color coded by chromosome and has a horizontal line representing the statistical cutoff (Figure 2). Another feature of the COMICS package is the ability to download the data frame generated from the ICS. Specifically, the output file contains six different columns, including the chromosome, the SNP position (or midpoint position in a window), outlier status (Boolean form), ICS Distance, logged ICS, and the position with respect to the chromosome. Importantly, the downloadable content in COMICS allows for the flexibility of both downstream analyses of multi-dimensional data and novel figure generation.

**Figure 1.**
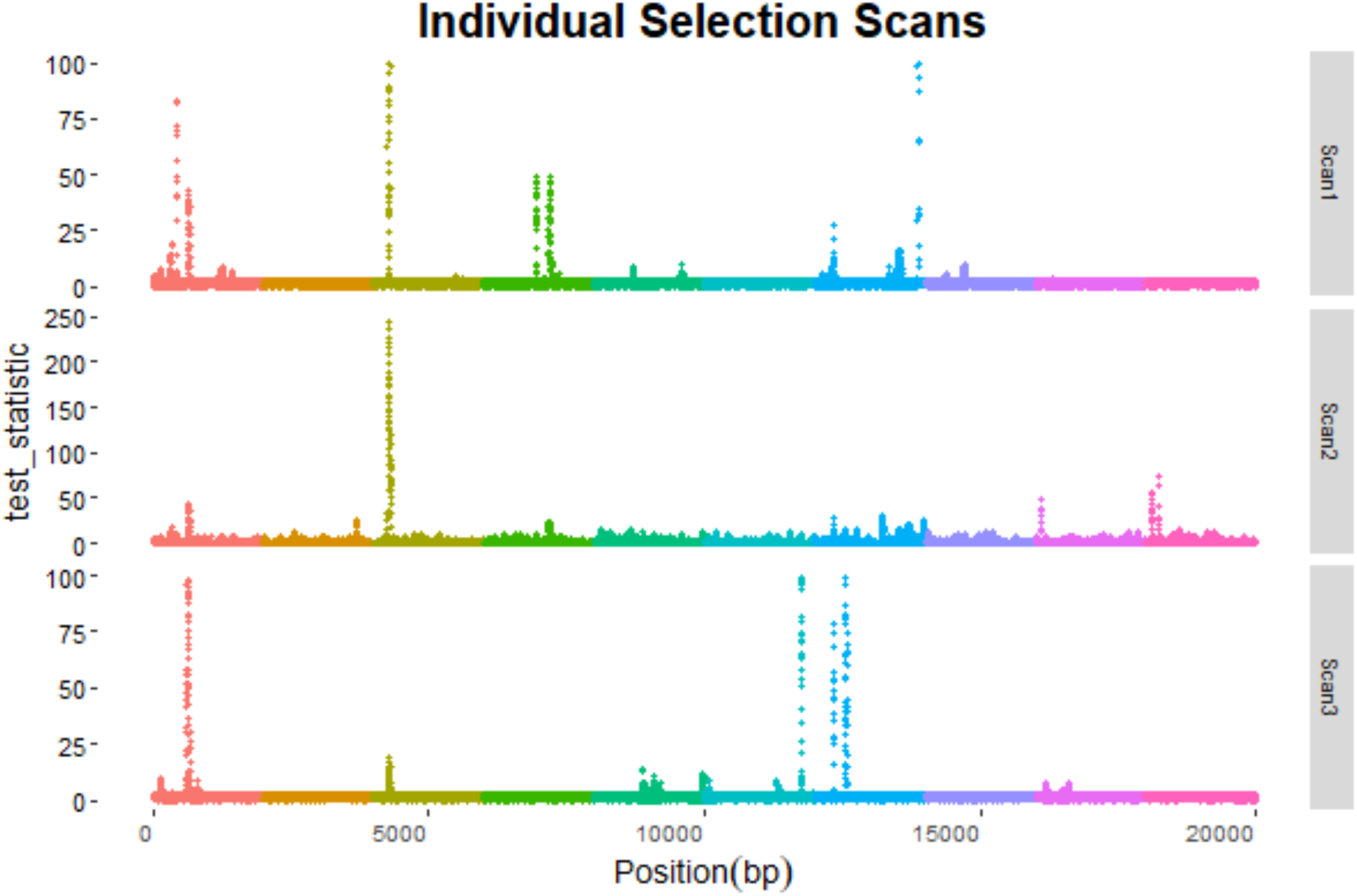
A screenshot of the suite of simulated selection scans that can be generated with COMICS. COMICS also allows the user to create individual selection scans of each univariate statistic that contributes the multidimensional scatter. In this example, there are three different selection methods that contribute to the ICS distance, Scan1, Scan2, and Scan3. Note that each color change represents a different chromosome. For simplicity, this example figure contains only ten 2kb chromosomes.

**Figure 2.**
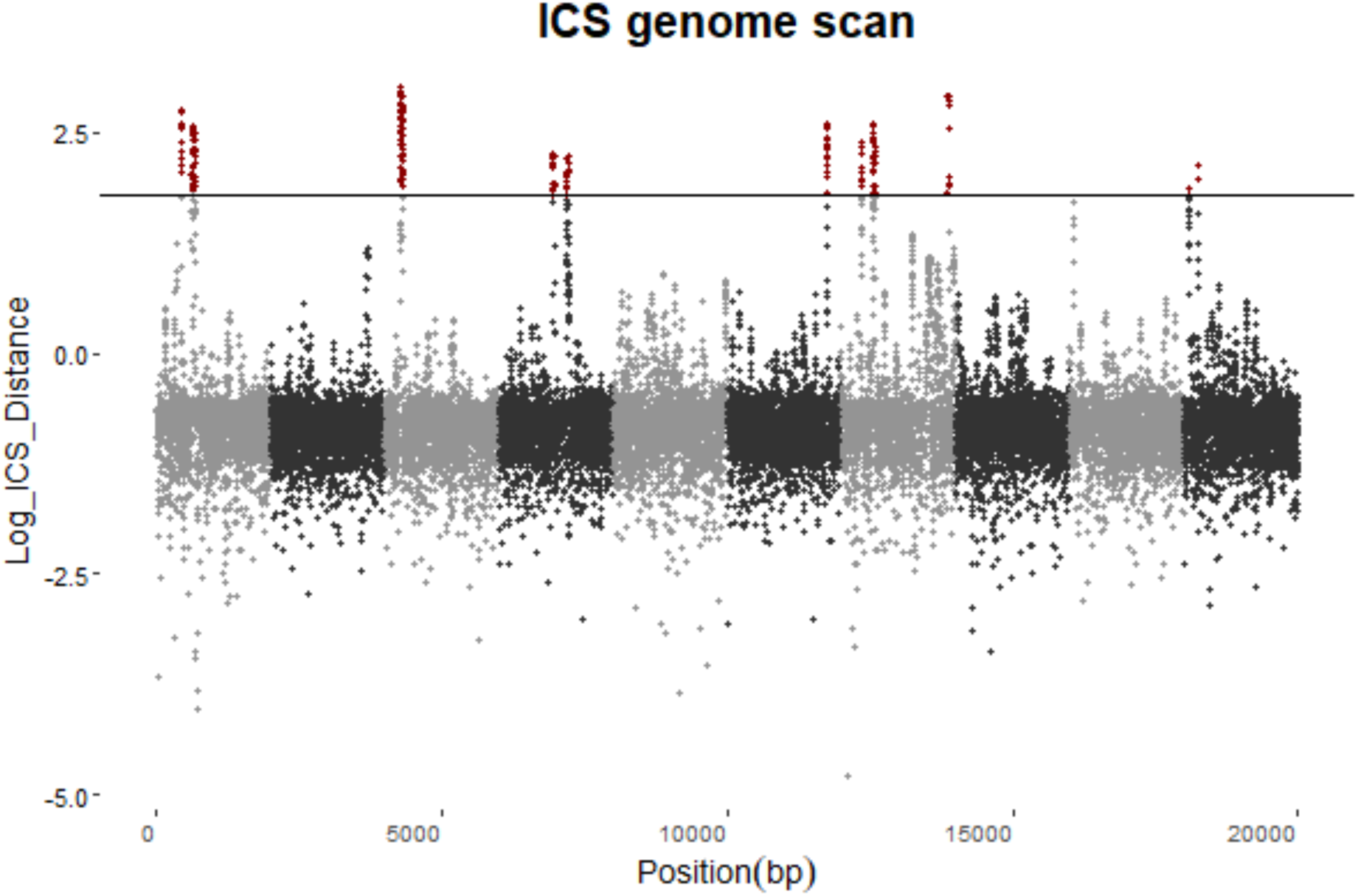
A screen shot from the COMICS GUI showing the final genome scan once transformed to the multi-dimensional scatter. The Y-axis is the ICS distance while the X-axis is the position along the genome. The color difference between grey and black represents transitions from one chromosome to another. The black horizontal line is the statistical cutoff that is defined by the user. Depending on the statistical cutoff, COMICS will render all data points above the threshold red where all other loci will conform to the default chromosome color. Note that the chromosomes numbers and sizes are the same as Figure 1.

### Testing the ICS with genomic data

Since the development of the ICS, studies have shown that it is a robust method for detecting outliers in a multidimensional space (Tyler et al. 2007; Archimbaud, Aurore, Nordhausen, Ruiz-Gazen 2018a). However, none of these analyses have been geared toward genomic data. To test the robustness of the ICS, we used simulated genomic data from previously published analyses (Lotterhos and Whitlock 2014; Lotterhos and Whitlock 2015; Verity et al. 2017). Specifically,, genetic data were simulated under several demographic models and include populations that experience either an equilibrium or non-equilibrium state. For example, we used data simulated under four different demographic scenarios: a non-equilibrium isolation by distance due to a one refugium and two refugia demographic expansion, an equilibrium isolation by distance model where the landscape was started at carrying capacity and was run until equilibrium, and a island model at equilibrium with a migration rate of 0.01 among demes. All simulated scenarios contained 30 populations that consisted of 20 individuals. The simulated data also consisted of 10,000 independent SNPs where 9,900 SNPs were evolving neutrally and the remaining 100 SNPs were under the influence of natural selection with different selection coefficients: 12 SNPs where s = 0.1, 38 SNPs where s = 0.01, and 50 SNPs where s = 0.005. Specifically, the strength of selection within a simulated population was determined by the standardized environment for that specific population; if observed environmental values were greater than the mean environment then selection was positive, however neutrality was concluded if there were differences between observed environmental values and the average environment (cite). For each SNP, five different univariant statistics were used that were calculated from previously published studies (cite sim studies). These statistics include 1) log-Bayes factor (a measure of the association between allele frequency and the environment calculated in BAYENV2, 2) Spearman’s rho (additional measure of association between alleles and the environment from BAYENV2, 3) X^T^X (measure of genetic differentiation among populations from BAYENV2, 4) Z-score (a measure of association between genotypes and the environment from the LFMM R-package),(Gunther and Coop 2013) (cite LFMM). For a more comprehensive description of the previously simulated data see (Lotterhos and Whitlock 2014; Lotterhos and Whitlock 2015).

To directly compare between Mahalanobis and ICS, we identified the number of true-positives/negatives, false-positives/negatives, and recall (the number of true-positives over the number of putative positives identified) as a measure of performance. We display this information in the form of a confusion matrix (Figure 3B & 3D). A confusion matrix is a table that is used to describe the performance of a defined model (ICS and Mahalanobis distances) on a test data set of known values (simulated data). In our confusion matrices, there are four different categories that are defined by a Boolean format: Actual Neutral, Identified Neural, Actual Selection, and Identified Selection; where “Actual” is defined by the known values from the simulated data and “Identified” is defined by the outputs of the ICS and Mahalanobis. For example, the number of loci that were “Actual Neutral” and “Identified Neutral” are true-negatives and those that were “Actual Selection” and “Identified Selection” are true-positives.

**Figure 3.**
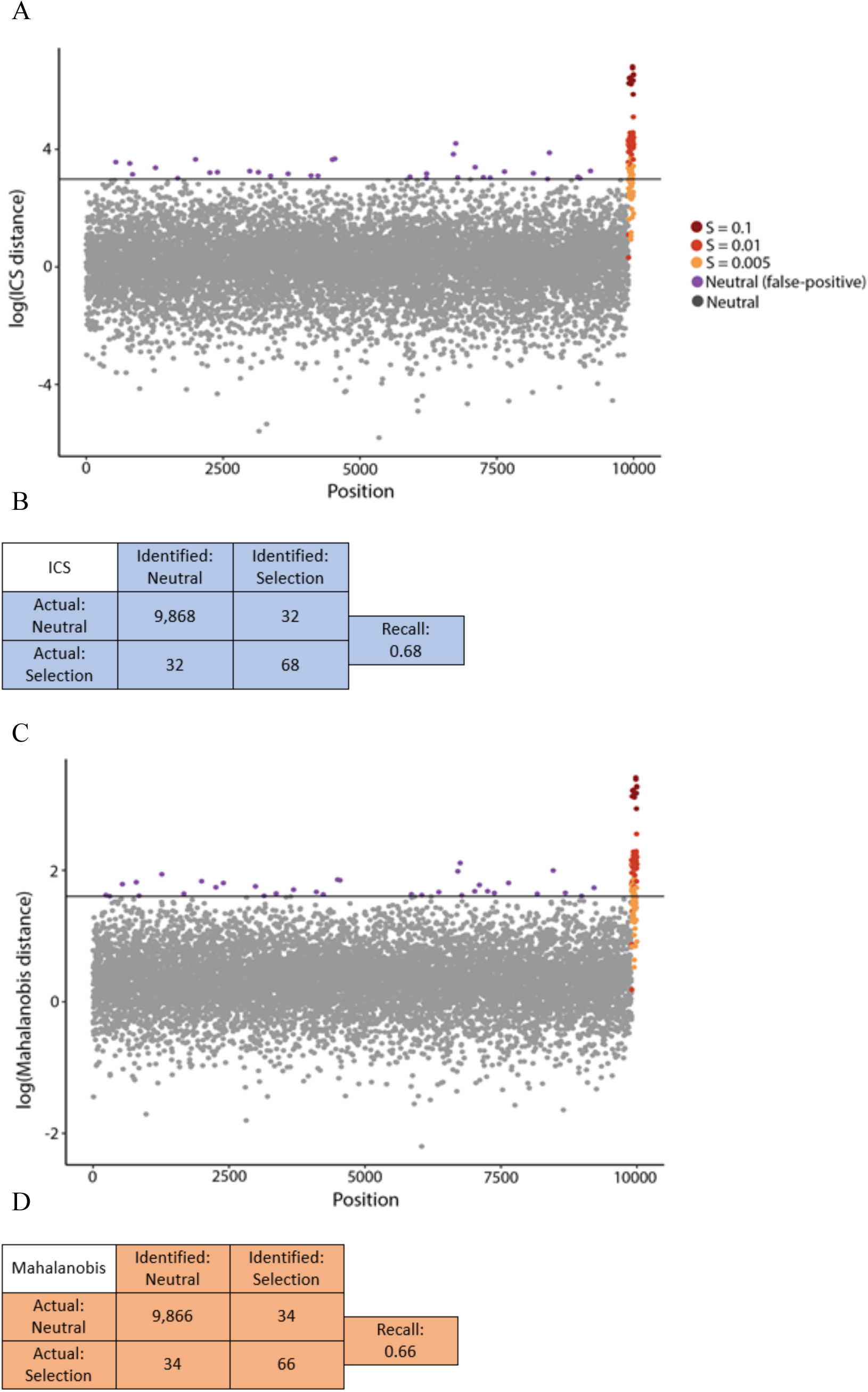
Selection scans and confusion matrices for the ICS (A & B) and Mahalanobis (C &D) distances calculated using simulated data. The horizontal line represents our statistical cutoff for outlier loci where all true-negatives are marked as grey and false-positives are purple. Loci that were simulated with a selection coefficient are in shades of red where the darkest represent loci with the largest selection coefficient. Importantly, loci with a selection coefficient that were below the statistical cutoff were deemed false-negatives and those above the line were true-positives. Within confusion matrices, the sum of the columns and the sum of the rows equals the total number of loci used in the simulated data.

In the COMICS application, outliers are determined by a statistical cutoff. Statistical cutoffs in COMICS are defined by the user and are picked out of the distribution of ICS distances, usually in the form of quantiles (i.e., top 1% of ICS distances). Importantly, the detection of outliers only satisfies a statistical pattern which may or may not be indicative of the biological process of selection. Because we are using simulated data we are able to make a clear distinction between outliers that are the result of a statistical pattern (i.e., arbitrary statistical cutoffs) versus those that are identified as an outlier from a biological process (i.e., natural selection). For our comparative analyses we used a 1% cutoff. Of the 10,000 simulated loci (9,900 neutral and 100 selected). Moreover, the ICS performed marginally better than Mahalanobis distance when separating selected loci from the neutral background across loci under varying selective coefficients. (Figure 3A). Out of the top 1% (the first 100 loci with the largest ICS distance), there was a total of 32 false-positives, with the remaining 68 loci being true-positives. In terms of neutral loci, the ICS identified a total of 9,868 loci as true-negatives with 32 false-negatives (Figure 3B). Across different regimes of selective coefficients, all loci with a large selective coefficient (s = 0.1) were identified as an outlier, 94.7% (35 of 38) of loci with intermediate selection coefficient (s = 0.01) were correctly identified as an outlier, while 42% of weakly selected loci (s = 0.005) were identified as outliers. The fraction of relevant outliers versusthe total of relevant outliers (recall) for ICS was 0.68; meaning that out of the top 100 ICS distances, 68% of loci were under selection (Figure 3A). Analysis with Mahalanobis distances allowed us to correctly identify loci with strong selection coefficients (s = 0.1) as outliers. 94.7% of loci with intermediate coefficients (s = 0.01) were correctly identified as outliers, and only 38% of weakly selected (s = 0.005) loci were correctly identified as outliers (Figure 3C). The total recall for selected loci under the Mahalanobis distance was 0.66 (Figure 3D). When ICS results are compared with those obtained with Mahalanobis distances, we found nearly a threefold decrease in false-positives (total false-positives in Mahalanobis = 34, as opposed to 32 in ICS) as well as an increase in the true-positives (true-positives in Mahalanobis = 66 as opposed to 68 in ICS, Figure 3B & 3D).

## Discussion

As the amount of genetic data increase, so does the number of SNPs (or windows) that can be tested for neutral or selective evolution. Because each SNP (or window) is considered an individual hypothesis, confounding effects on genetic analyses including the inflation of false-positives and false-negatives will become more prominent. The inflation of type I and type II errors brings further issues pertaining to the interpretation of putatively selected genetic markers and their implication on downstream analyses; for example, the misidentification of adaptive loci or the inability to capture adaptive loci. There is a growing need in developing methods to combine methods of selection, as the number of methods keeps growing in numbers. Here we developed, tested, and introduced an R-package (COMICS) using the Shiny smart wrapper that incorporates the use ICS specifically for genomic data.

We show that ICS is a robust method for identifying outlier loci in multi-dimensional space and propose its use over other similar existing methods like Mahalanobis distances. Specifically, the ICS identified fewer false-positives/negatives, more true-positives/negatives and had a higher recall for true-positives. One of the more interesting aspects of the ICS was its ability to detect a larger proportion of weakly selected loci. Across the genome and across a geographic space, loci experience selective pressures at varying strengths. Across methods, signatures of loci under strong selection are often identified as true-positives. As the strength of selection acting on a locus is reduced, the chances of identifying it as a true outlier are also reduced, even after combining multiple selection detection methods that vary in their ability to detect signatures of selection. Compared to contemporary methods, the ICS offers an increase in the recall for outlier loci even for weakly selected loci. We therefore recommend that future studies using univariate statistics to identify loci under selection consider the ICS as means to identify outliers in multi-dimensional space.

## References

Alachiotis N, Stamatakis A, Pavlidis P. 2012. OmegaPlus: a scalable tool for rapid detection of selective sweeps in whole-genome datasets. Bioinformatics. 28:2274–2275.

Anopheles gambiae 1000 Genomes Consortium. 2017. Genetic diversity of the African malaria vector Anopheles gambiae. Nature. 552:96.

Archimbaud A, Nordhausen K, Ruiz-Gazen A. 2016. ICSOutlier: Outlier Detection Using Invariant Coordinate Selection. R Package Version 0.2-0. URL Http://CRAN.R-Project.Org/package=ICSOutlier.

Archimbaud A, Nordhausen K, Ruiz-Gazen A. 2018a. ICS for multivariate outlier detection with application to quality control. Comput Stat Data Anal. 128:184–199.

Archimbaud A, Nordhausen K, Ruiz-Gazen A. 2018b. ICSOutlier: Unsupervised Outlier Detection for Low-Dimensional Contamination Structure. R Journal. 10.

Beaumont MA, Balding DJ. 2004. Identifying adaptive genetic divergence among populations from genome scans. Mol Ecol. 13:969–980.

Bekkevold D, Gross R, Arula T, Helyar SJ, Ojaveer H. 2016. Outlier loci detect intraspecific biodiversity amongst spring and autumn spawning herring across local scales. PloS One. 11:e0148499.

Bellman R. 2013. Dynamic programming. New York: Courier Corporation.

Bellman RE. 2015. Adaptive control processes: a guided tour. Princeton university press.

Campbell‐Staton S, Edwards S, Losos J. 2016. Climate‐mediated adaptation after mainland colonization of an ancestrally subtropical island lizard, A nolis carolinensis. J Evol Biol. 29:2168–2180.

Capblancq T, Luu K, Blum MG, Bazin E. 2018. Evaluation of redundancy analysis to identify signatures of local adaptation. Molecular Ecology Resources. 18:1223–1233.

Chang W, Cheng J, Allaire JJ, Xie Y, McPherson J. 2015. Shiny: web application framework for R. R Package Version 0.11. 1:106.

Chen H, Patterson N, Reich D. 2010. Population differentiation as a test for selective sweeps. Genome Res. 20:393–402.

De Maesschalck R, Jouan-Rimbaud D, Massart DL. 2000. The mahalanobis distance. Chemometrics Intellig Lab Syst. 50:1–18.

Fischer D, Honkatukia M, Tuiskula-Haavisto M, Nordhausen K, Cavero D, Preisinger R, Vilkki J. 2017. Subgroup detection in genotype data using invariant coordinate selection. BMC Bioinformatics. 18:173.

Forester BR, Jones MR, Joost S, Landguth EL, Lasky JR. 2016. Detecting spatial genetic signatures of local adaptation in heterogeneous landscapes. Mol Ecol. 25:104–120.

Gnanadesikan R, Kettenring JR. 1972. Robust estimates, residuals, and outlier detection with multiresponse data. Biometrics.:81–124.

Gunther T, Coop G. 2013. Robust identification of local adaptation from allele frequencies. Genetics. 195:205–220.

Hodel RG, Chandler LM, Fahrenkrog AM, Kirst M, Gitzendanner MA, Soltis DE, Soltis PS. 2018. Linking genome signatures of selection and adaptation in non-model plants: exploring potential and limitations in the angiosperm Amborella. Curr Opin Plant Biol. 42:81–89.

Leys C, Klein O, Dominicy Y, Ley C. 2018. Detecting multivariate outliers: Use a robust variant of the Mahalanobis distance. J Exp Soc Psychol. 74:150–156.

Lotterhos KE, Whitlock MC. 2015. The relative power of genome scans to detect local adaptation depends on sampling design and statistical method. Mol Ecol. 24:1031–1046.

Lotterhos KE, Whitlock MC. 2014. Evaluation of demographic history and neutral parameterization on the performance of FST outlier tests. Mol Ecol. 23:2178–2192.

Mahalanobis PC. 1936. On the generalized distance in statistics. 26:541–588.

Martins H, Caye K, Luu K, Blum MG, Francois O. 2016. Identifying outlier loci in admixed and in continuous populations using ancestral population differentiation statistics. Mol Ecol. 25:5029–5042.

Nielsen R, Williamson S, Kim Y, Hubisz MJ, Clark AG, Bustamante C. 2005. Genomic scans for selective sweeps using SNP data. Genome Res. 15:1566–1575.

Pavlidis P, Živković D, Stamatakis A, Alachiotis N. 2013. SweeD: likelihood-based detection of selective sweeps in thousands of genomes. Mol Biol Evol. 30:2224–2234.

Rellstab C, Zoller S, Walthert L, Lesur I, Pluess AR, Graf R, Bodénès C, Sperisen C, Kremer A, Gugerli F. 2016. Signatures of local adaptation in candidate genes of oaks (Quercus spp.) with respect to present and future climatic conditions. Mol Ecol. 25:5907–5924.

Sabeti PC, Varilly P, Fry B, Lohmueller J, Hostetter E, Cotsapas C, Xie X, Byrne EH, McCarroll SA, Gaudet R. 2007. Genome-wide detection and characterization of positive selection in human populations. Nature. 449:913.

Sabeti PC, Reich DE, Higgins JM, Levine HZ, Richter DJ, Schaffner SF, Gabriel SB, Platko JV, Patterson NJ, McDonald GJ. 2002. Detecting recent positive selection in the human genome from haplotype structure. Nature. 419:832.

Team RC. 2018. R: A Language and Environment for Statistical Computing.Vienna: R Foundation for Statistical Computing; 2018. ISBN:241–262.

Trunk GV. 1979. A problem of dimensionality: A simple example. IEEE Trans Pattern Anal Mach Intell.:306–307.

Tyler D, Critchley F, Dümbgen L, Oja H. 2007. Invariant coordinate selection. Conditionally Accepted.

Verity R, Collins C, Card DC, Schaal SM, Wang L, Lotterhos KE. 2017. minotaur: A platform for the analysis and visualization of multivariate results from genome scans with R Shiny. Molecular Ecology Resources. 17:33–43.

Wickham H. 2016. ggplot2: elegant graphics for data analysis. Springer.

Wickham H. 2007. Reshaping data with the reshape package. Journal of Statistical Software. 21:1–20.

Zhong M, Zhang Y, Lange K, Fan R. 2011. A cross-population extended haplotype-based homozygosity score test to detect positive selection in genome-wide scans. Statistics and its Interface. 4:51.

Zueva KJ, Lumme J, Veselov AE, Kent MP, Primmer CR. 2018. Genomic signatures of parasite-driven natural selection in north European Atlantic salmon (Salmo salar). Marine Genomics. 39:26–38.

